# Performance of *Orbicella faveolata* larval cohorts does not align with previously observed thermal tolerance of adult source populations

**DOI:** 10.1101/2023.06.28.546780

**Authors:** Yingqi Zhang, Shelby E. Gantt, Elise F. Keister, Holland Elder, Graham Kolodziej, Catalina Aguilar, Michael S. Studivan, Dana E. Williams, Dustin W. Kemp, Derek P. Manzello, Ian C. Enochs, Carly D. Kenkel

## Abstract

*Orbicella faveolata*, commonly known as the mountainous star coral, is a dominant reef-building species in the Caribbean, but populations have suffered sharp declines since the 1980s due to repeated bleaching and disease-driven mortality. Prior research has shown that inshore adult *O. faveolata* populations in the Florida Keys are able to maintain high coral cover and recover from bleaching faster than their offshore counterparts. However, whether this origin-specific variation in thermal resistance is heritable remains unclear. To address this knowledge gap, we produced purebred and hybrid larval crosses from *O. faveolata* gametes collected at two distinct reefs in the Upper Florida Keys, a nearshore site (Cheeca Rocks, CR) and an offshore site (Horseshoe Reef, HR), in two different years (2019, 2021). We then subjected these aposymbiotic larvae to severe (36 °C) and moderate (32 °C) heat challenges to quantify their thermal tolerance. Contrary to our expectation based on patterns of adult thermal tolerance, HR purebred larvae survived better and exhibited gene expression profiles that were less driven by stress response under elevated temperature compared to purebred CR and hybrid larvae. One potential explanation could be compromised reproductive output of CR adult colonies due to repeated summer bleaching events in 2018 and 2019, as gametes originating from CR in 2019 contained less storage lipids than those from HR. These findings provide an important counter-example to the current selective breeding paradigm, that more tolerant parents will yield more tolerant offspring, and highlight the importance of adopting a holistic approach when evaluating larval quality for conservation and restoration purposes.

## Introduction

Global ecosystems are undergoing unprecedented structural and functional changes as atmospheric CO_2_ level and temperature continue to rise in the Anthropocene (Steffen et al., 2007). One ecosystem that is particularly vulnerable to these changes is coral reefs, because most reef-building corals are found in the tropics (Spalding & Brown, 2015) and already live close to their upper thermal limits (Baker et al., 2008). A small temperature increase, as little as 1 °C above the maximum monthly mean temperature for a period of four weeks, or four degree heating weeks (Liu et al., 2005), can lead to the breakdown of the symbiotic relationship between the cnidarian animal host and their intracellular photosynthetic dinoflagellate algae. This phenomenon is commonly known as coral bleaching (Hoegh-Guldberg et al., 2007; Lesser, 2011). Worldwide, coral cover is estimated to have declined by 20% over the past 30 years and reefs will continue to be threatened by large-scale bleaching events even with climate intervention strategies (Hoegh-Guldberg et al., 2019). Similar to the pattern observed in the wider Caribbean region (Gardner et al., 2003), coral reefs in the Florida Keys have experienced drastic population declines since the early 1980s mostly due to bleaching and disease (Dustan & Halas, 1987; Precht & Miller, 2007). The two most recent large-scale bleaching events to affect this region occurred in 2014 and 2015 when maximum temperatures exceeded local bleaching thresholds for over 4-8 weeks (Eakin et al., 2019; Smith et al., 2019).

However, not all corals are equally susceptible to bleaching. Coral populations inhabiting thermally-challenging environments, characterized by elevated temperatures and/or greater temperature variabilities, have been repeatedly shown to exhibit higher tolerance to heat stress (Howells et al., 2016; Thomas et al., 2018). Mechanistically, increased temperature tolerance can be the result of adaptation and/or acclimatization on the part of coral hosts, their dinoflagellate endosymbionts, or other members of the microbiome (Ainsworth et al., 2016; Berkelmans & van Oppen, 2006; Palumbi et al., 2014; Santoro et al., 2021). Along the Florida

Keys reef tract, inshore patch reefs experience higher annual temperature fluctuations and elevated mean temperature during bleaching-prone summer months in comparison to offshore reefs at similar latitudes (Kenkel et al., 2015; Manzello et al., 2015a, 2015b). This spatially defined thermal heterogeneity has been theorized to support elevated heat tolerance of inshore corals, which aligns with lab-based experiments and field-based observations of reduced bleaching severity of inshore coral populations (Gintert et al., 2018; Kenkel et al., 2013). During the back-to-back bleaching events in 2014 and 2015, inshore *Orbicella faveolata* colonies in the Upper and Lower Florida Keys demonstrated lower bleaching prevalence and higher recovery rate than colonies at paired offshore sites (Manzello et al., 2019). Due to the lack of distinct genetic structure among inshore and offshore host populations, the increased heat tolerance of the inshore corals was attributed to the significantly greater prevalence of heat tolerant symbionts (*Durusdinium trenchii*) in these corals versus those at offshore sites (Manzello et al., 2019). The host role in shaping holobiont thermotolerance in this system remains unclear.

Establishing the degree to which the coral host contributes to thermal tolerance is essential for modeling adaptive potential and implementing intervention strategies. Assisted evolution was proposed as a suite of intervention approaches to mitigate the decline and degradation of reef systems, given that adaptive changes that occur naturally might not be able to keep pace with the rapidly-changing climate (Van Oppen & Oliver, 2015). Studies in multiple Indo-Pacific coral species provide compelling evidence for host genomic heritability of traits to enhance the tolerance to heat, ocean acidification, and disease (Dixon et al., 2015; Drury et al., 2022; Howells et al., 2021; Quigley et al., 2020). Additional data is needed to better understand the tradeoffs of selecting for a single trait in corals, given their exposure to multiple environmental challenges (Ladd et al., 2017). This approach may also lead to outbreeding depression and genetic swamping, which can threaten the survival and fitness of their offspring (Aitken & Whitlock, 2013). Significant differences in the decline of coral species have been observed in the Pacific and Caribbean regions (Tebbett et al., 2023)). It has been suggested that impaired colony physiology in corals could contribute to suboptimal larval performance that causes recruitment failure (Hughes & Tanner, 2000; Williams et al., 2008).

We investigate how the source population affects the performance of *O. faveolata* offspring. To do this, we created larval cohorts sourced from parent colonies living in thermally distinct reef sites in the Upper Florida Keys. We then exposed these aposymbiotic larvae to severe and moderate temperature stress.*O. faveolata* is one of the major reef-building corals in the Florida Keys and its populations are highly connected throughout the wider Caribbean seascape ((Rippe et al., 2017)). However, *O. faveolata* populations have suffered sharp declines in the past few decades (Edmunds, 2015). These declines are largely due to bleaching and disease, making recovery challenging (Gladfelter et al., 1978). *Orbicella faveolata* is a hermaphroditic broadcast-spawning coral species that sexually reproduces during late summer months when water temperatures are maximal (Szmant, 1991). Symbiotic dinoflagellates are acquired from the environment during metamorphosis (Coffroth et al., 2001). By working with aposymbiotic larvae, we can study the physiological and transcriptomic basis for heat tolerance in the animal host without the confounding effects of symbiosis. Understanding the physiological and genetic factors underlying origin-dependent bleaching resistance in *O. faveolata* (and congeners) is crucial because they were listed as threatened under the Endangered Species Act in 2014. By studying this, we can assess the impact on future generations and estimate adaptive capacity, enabling informed conservation efforts.

## Methods

### Sites and Temperature Data

Two well-monitored sites in the Upper Florida Keys were chosen for subsequent spawning collections in 2019 and 2021, including Cheeca Rocks (CR, 24.8977°N, 80.6182°W) and Horseshoe Reef (HR, 25.1388°N, 80.3133°W). Temperature was measured every 3 hrs at the Cheeca Rocks Moored-Autonomous pCO_2_ buoy (MApCO_2_, depth = 1 m) using a conductivity– temperature sensor (Model SBE-16 plus v. 2.2, Seabird Electronics). Data were collected every 30 min from HR (depth = 3.4 m) using the following loggers: Pendant (from 1/1/2017 to 8/6/2019 09:00 h) and Tidbit MX2204 (8/6/19 09:30 h to 12/31/21). From these data, daily average temperature and the running 30-day mean temperatures were calculated for each site from January 1, 2017 to December 31, 2021.

The bleaching thresholds for CR were determined as previously described (Gintert et al., 2018). Bleaching thresholds can be estimated for reef sites in the Florida Keys by taking the average of the maximum monthly mean sea surface temperature (SST) during a non-bleaching year and the minimum monthly mean SST during a bleaching year (Manzello et al., 2007). Every Florida Keys-wide mass bleaching event since 2005 has been predicted in near-real-time by calculating the running 30-day mean SST from the Molasses Reef Coastal Marine Automated Network (C-MAN) station and using that site as a proxy for the rest of the Florida Keys offshore reef sites (Manzello, 2015). This technique has also been used to predict every bleaching event that has occurred at CR since 2012.

The bleaching threshold for CR is a monthly mean SST ≥ 31.3°C, such that once the running 30-day mean SST reach this value, bleaching has been observed in 2014, 2015, 2018, and 2019 (Gintert et al., 2018; Manzello et al., 2019) (Fig. 1). The monthly mean bleaching threshold for offshore reefs in the Florida Keys is ≥ 30.4°C, nearly 1°C lower than CR. We lack sufficient coverage of both bleaching and non-bleaching year temperature data for HR to determine a local bleaching threshold for this site. However, we can deduce that temperatures experienced at HR are warmer than the values experienced at the far offshore reefs used for calculation of the Florida Keys-wide threshold, but cooler than CR (Fig. 1).

**Fig. 1.**
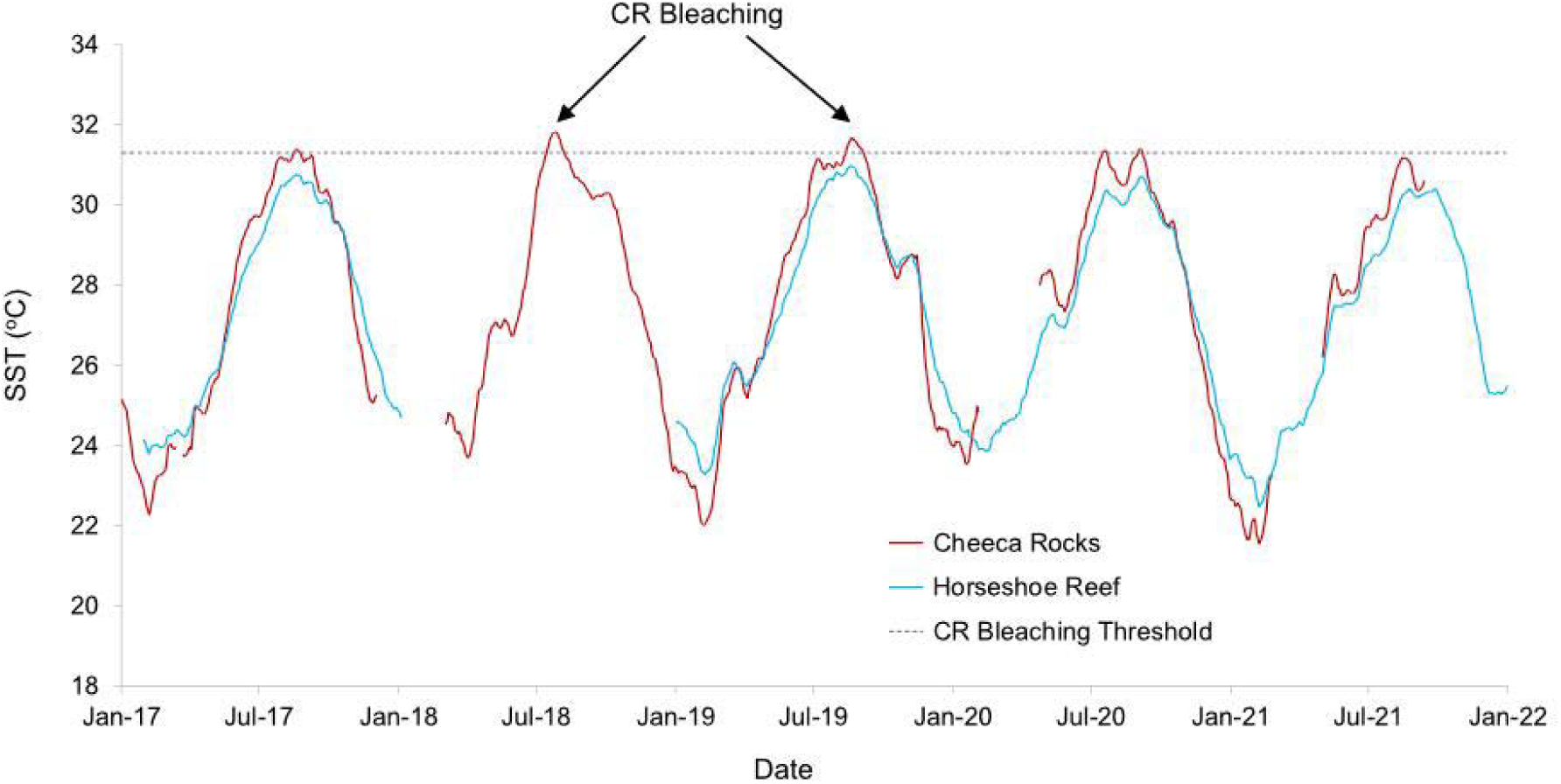
Time series of running 30-day mean sea surface temperature (SST) for Cheeca Rocks (red line) and Horseshoe Reef (blue line) from 2017 to 2021. Bleaching threshold for Cheeca Rocks (monthly mean SST ≥ 31.3°C) shown as dashed black line.

### Spawning, cross design and larval rearing

Spawning collections were conducted under permit FKNMS-2018-163-A1. The first spawn occurred on August 22, 2019, seven days after the full moon. Gamete bundles were collected from five adult colonies within each site using spawning tents with 50 mL collection tubes following standard protocols (Marhaver et al. 2017). Collection tubes were removed from tents after ∼5 mL of gametes had been collected, immediately capped, and transported back to the boat by divers. Gametes were then diluted to reach a sperm concentration ofL∼L10^6^ cells/mL in individual 5 gallon buckets filled with 0.2 μm filtered seawater (FSW). Three replicate bulk crosses were created on each boat by mixing equivalent aliquots of diluted gametes from each colony (n = 5) at each site (n = 2). Approximately 1.5 hrs post spawning, diluted gametes released from two CR colonies and three HR colonies were mixed in the laboratory to create three replicate hybrid bulk crosses. Note that given the optimum fertilization window for gametes (∼2 hrs), there was not sufficient time to separate eggs from sperm to attempt a diallel-type crossing design. Therefore, the bulk hybrid crosses could include true CR × HR and HR × CR reciprocal larvae as well as CR × CR and HR × HR fertilizations.

A second round of *O. faveolata* gamete collection from each site occurred six (28 August, HR only) and seven (29 August, CR and HR) days after the full moon in August 2021. Gametes were again obtained from five colonies at HR on day 6, and handled as in 2019, resulting in three bulk crosses of HR × HR cultures. No spawning was observed at CR on day 6. On day 7, only a single colony was observed spawning at each site, which restricted our fertilization design to the creation of hybrid crosses only. Gamete bundles from each site were returned to the Key Largo Marine Research Laboratory and separated into eggs and sperm by filtration through an 80 μm nitex mesh, followed by rinsing with FSW. Individual eggs and sperm were crossed to create three culture replicates of each of two hybrid crosses (CR sperm × HR egg, HR sperm × CR egg).

For all crosses in each year, successful fertilization was confirmed through observation of initial cell division under 100x magnification following ∼2 hrs of incubation. Developing embryos were gently rinsed 3x in FSW to remove excess sperm and transferred to 6 L culture bins at a density of ∼1 embryo per mL in FSW. Healthy developing larvae were rinsed and transferred to fresh FSW twice daily until reaching the planula stage, after which water changes were performed every other day. Larval cultures were maintained at 29°C by placing filled bins in shallow 32 L polycarbonate (Rubbermaid) water baths equipped with 100 W aquarium heaters and SL381 submersible pumps (Domica) to maintain ambient temperature consistent with field temperature profiles.

### Thermal stress challenges

Two thermal stress experiments were conducted at the Key Largo Marine Research Laboratory in 2019 after mature swimming larvae were observed in all cultures (CRxCR, HRxHR, putative hybrid cross), which occurred on day 3 post fertilization: an acute stress at 36 °C and a moderate stress at 32 °C following (Zhang et al., 2022). For each experiment, six replicate 6 L polycarbonate larval bins (Vigors) were filled with 0.2 μm FSW and placed into a set of two shallow 32 L polycarbonate (Rubbermaid) water baths for temperature control (n = 3 larval bins per bath). Each water bath was filled with ∼15 L water and equipped with a SL381 submersible water pump to maintain circulation and a 100 W aquarium heater. Each larval rearing bin was fully filled and two of three bins were equipped with HOBO temperature loggers (Onset). For the acute stress experiment, each larval rearing bin received two groups of ten larvae per bulk cross (n = 6 per cross type per treatment) that were aliquoted into floating netwells (70 μm cell strainers, Grenier Bio-One). The control bath remained at 29 °C and the treatment bath was heated up to 36 °C over 24 hrs (Fig. S1a). Mortality was assessed every 24 hrs by counting the number of surviving larvae in the netwells. The experiment was terminated once mortality reached more than 50% for the majority of the netwells. For the moderate stress experiment, a total of 20 larvae from each bulk cross replicate were aliquoted to each netwell. The control bin remained at 29 °C and the treatment bin was heated up to 32 °C over 24 hrs (Fig. S1a). After 4 days of exposure, swimming larvae were removed from each netwell, flash frozen in liquid nitrogen, and stored at -80 °C for RNA extraction.

Similar acute and moderate stress experiments were conducted with larvae reared in 2021 (HR × HR, CR × HR, HR × CR). All cultures were transported to the Experimental Reef Lab (ERL) at The University of Miami’s Cooperative Institute for Marine and Atmospheric Studies (CIMAS) on day 4 post fertilization. Larvae were packed into 50 mL centrifuge tubes with no air bubbles and stored in coolers at ambient temperature during transit. Upon arrival at ERL, larvae were re distributed into 6 L culture bins (n = 3 per cross) filled with 0.2 μm FSW. Temperature control was accomplished using the ERL aquaria with individual treatment tanks serving as water baths (Enochs et al., 2018). For the stress experiments, netwells were floated directly in the temperature-controlled flow-through aquaria. For each bulk cross, two groups of 10 larvae were allocated to each treatment tank in the acute stress treatment (n = 6 per cross type per treatment) and three groups of 20 larvae were allocated to each treatment tank in the moderate stress (n = 9 per cross type per treatment). The control temperature for both experiments was set to 27 °C. Heat ramps started 5 days post fertilization (note this represented different calendar days for HR × HR vs. CR × HR and HR × CR to account for differences in developmental age) and target temperatures were reached over 48 hrs (Fig. S1b). Lights were maintained at 180 µmol s^-1^·m^-2^ for a 12:12 hr light dark cycle. The acute stress assay was monitored every 12 hrs for survival and the experiment was terminated once mortality reached more than 50% for the majority of the netwells. For the moderate duration experiment, larvae were retrieved from 1 netwell per cross per replicate tank following 4 days of exposure using a pipette, counted, and transferred to a cryovial for RNA extraction. Excess seawater was removed and larvae were snap frozen in liquid nitrogen and stored at -80 °C until processing. Additional replicate samples were taken from the remaining 2 netwells for protein and lipid analyses.

### Physiological assays

Gametes from both years, as well as 2021 fertilized larvae, were collected for lipid analyses. Gametes from each parent colony were sampled in duplicate. Gametes and larvae were counted under a dissection microscope before being transferred to combusted glass tubes and frozen at -20 °C until lipid processing. Total lipids were extracted and determined gravimetrically using a modified Folch method (Folch et al., 1957), as described in (Keister et al., 2023). Subsequently, 100% chloroform was added to total lipids to achieve a 10 mg mL^-1^ concentration, to standardize samples. Lipid classes were quantified by spotting, in duplicate, 1 μL of extracted lipids on silica Chromarods^®^ before being developed using thin-layer chromatography via a two-step solvent system (Conlan et al., 2014, 2017; Nichols et al., 2001), as described in (Keister et al., 2023). Developed rods were then dried at 100 °C for 10 min in an IsoTemp Oven (Fisher Scientific) before being run on an Iatroscan MK 6S thin-layer flame ionization detector (TLC-FID). Known concentrations of lipid compounds, ranging from 0.1–10.0 mg mL^-1^, were used to calibrate the Iatroscan for the following lipid classes: phosphatidylethanolamine (PE), phosphatidylserine and phosphatidylinositol (PS-PI), phosphatidylcholine (PC), lysophosphatidylcholine (LPC), wax esters (WAX), triacylglycerols (TAG), sterols (ST), and diacylglycerols (DAG). All phospholipid lipid classes (PE, PS-PI, PC, LPC) were grouped and analyzed as one unit. All total lipid and lipid class values were presented as μg per gamete or μg per larva, respectively.

Protein samples were thawed on ice and homogenized by back pipetting in FSW for at least 60 s until no visible cellular debris was present. Total homogenate volume was recorded. Soluble host protein was quantified in triplicate with a BCA Protein Assay Kit II (BioVision) following the manufacturer’s protocol. Final protein concentration was multiplied by the initial homogenate volume and then standardized by the number of total sampled larvae.

### Statistical analysis of physiological trait data and survival under acute stress

All statistical analyses were conducted in R 4.2.1 (R Core Team, 2022). Traits were evaluated for normality using the Shapiro-Wilks test and log-transformed if not normally distributed. A two-way anova was performed to analyze the effects of treatment (levels: control and heat) and origin (levels: CR × CR, HR × HR) on 2019 gamete lipid content. No statistical analysis of 2021 gametes was possible given that only one CR colony spawned. Larval protein and lipid content for the 2021 cohorts were analyzed using a two-way anova to analyze the effect of origin (levels: HR × HR, CR × HR, HR × CR) and treatment (levels: control and heat). A mixed effects Cox model (Therneau, 2018) was used to model time of death for the acute stress experiment from both years as a function of treatment and origin, including a random effect of replicate bulk cross. Due to low mortality rate, HR × HR and control were chosen as the reference level for each fixed effect to calculate hazard ratio for other levels. A hazard ratio above 1 means a higher mortality risk and below 1 means a lower mortality risk compared to the reference levels. The null hypothesis was rejected at an alpha of 0.05.

### RNA extraction, library preparation, and sequencing

Total RNA was extracted from frozen samples using the Aurum Total RNA Mini Kit (Bio-Rad). Samples were back pipetted to ensure complete homogenization after Lysis Solution was added. Genomic DNA was removed by adding DNAse I on-column according to the manufacturer’s instructions. RNA concentration was quantified using a Take2 plate on a

Synergy H1 microplate reader (Biotek) and only samples with > 10 ng/µl concentration were used to generate tag-based RNA-seq libraries, following protocols modified for sequencing on the Illumina platform https://github.com/ckenkel/tag-based_RNAseq.

Libraries from 2019 (n = 45 samples) were sequenced on the NextSeq 550 in 2020 by the USC Genome Core using a 1×75bp HO kit. Libraries from 2021 (n = 30 samples) were sequenced on the NextSeq 2000 in 2022 in two replicate separate runs by the USC Norris Comprehensive Cancer Center Molecular Genomics core using a NextSeq 2000 P2 Reagents v3 kit, after which reads were concatenated to reach a comparable sequencing depth. The average sequencing depth per sample for the two libraries was 5.9 M reads (± 0.2 M) and 5.4 M reads (± 0.3 M) for the 2019 and 2021 datasets, respectively.

### Bioinformatics pipelines

Downstream bioinformatic processing was performed on USC’s Center for Advanced Research Computing (CARC) following protocols described in https://github.com/ckenkel/tag-based_RNAseq. Briefly, a custom perl script was used to discard PCR duplicates and reads missing adapter sequences. Poly-A tails and adapter sequences were trimmed and only high quality reads (PHRED score ≥ 20 over 90% of the read) were retained. A total of 2.7 M (± 0.09 M) and 2.4 M (± 0.1 M) reads per sample remained for the 2019 and 2021 datasets respectively after quality filtering. Filtered reads were mapped to an adult *Orbicella faveolata* transcriptome (Supplemental Materials) using the gmapper command from SHRiMP2 (Rumble et al., 2009). Read counts were summed by isogroup using a custom perl script, resulting in a per sample mapped read average of 1.5 M (± 0.05 M) and 1.3 M (± 0.07 M), for the 2019 and 2021 datasets respectively.

### Statistical analysis of gene expression

Larval samples from the two years were first compiled, where isogroups shared between the two year cohorts that had fewer than 2 counts across 90% of the samples were discarded, resulting in a total of 28,675 high quality isogroups remaining. Counts were then transformed using the *rlog* function in DESeq2 (Love et al., 2014) and outlier samples were identified through a sample network with a standardized connectivity score of less than −2.5, based on which one sample from the 2019 dataset and one from the 2021 data set were filtered. The filtered dataset was then separated by year to form two subsets, on which principal component analysis (PCA) was applied to visualize global gene expression pattern for each larval cohort using the R package FactoMineR (Lê et al., 2008).

Gene co-expression analysis was conducted using the WGCNA package in R (Langfelder & Horvath, 2008) following the standard tutorial (Langfelder & Horvath 2016) and a pipeline for performing meta-analysis of two microarray datasets (Miller 2011). Briefly, a signed gene co-expression network was constructed with a soft power of 4. Module assignment was generated based on the 2019 dataset and imposed onto the 2021 dataset to assess how well those modules were preserved across datasets. The eigengenes of these shared modules were correlated with origin (levels for 2019: CR × CR, HR × HR, Hybrid; levels for 2021: HR × HR, CR × HR, HR × CR) and treatment (levels: control, heat).

DESeq2 (Love et al., 2014) was used to examine genes that were differentially expressed in larvae by origin and treatment, with each year’s cohort being analyzed separately. Isogroups that had fewer than 2 counts across 90% of the samples were discarded, resulting in a total of 27,894 high quality reads in the 2019 dataset and 28,960 reads in the 2021 dataset. A two factor grouping was created by combining origin and treatment and fed into the DESeq2 model as the design, from which results were extracted with specific contrasts to obtain differentially expressed genes (DEGs) in response to treatment in a particular larval origin (e.g. CR × CR heat vs. CR × CR control). Significance testing was determined using a Wald test after independent filtering using a false discovery rate-corrected (FDR) threshold of 0.1. Multiple test correction was applied to raw p-values following Benjamini & Hochberg (Benjamini & Hochberg, 1995) and adjusted p-values less than 0.1 were considered significant. Signed log-p values were generated based on the adjusted p-values to serve as input for gene ontology (GO) enrichment analysis described below.

To explore the possibility that heat-responsive genes were front- or back-loaded in the CR × CR or HR × HR populations which would preclude their identification as significantly differentially expressed genes (Barshis et al., 2013), we identified the overlap between significantly up-regulated/down-regulated genes in response to heat in one population and the significantly up-regulated/down-regulated genes between groups in control conditions (i.e. constitutively differentially expressed) as candidate front-/back-loaded genes in the other population. For example, genes identified as significantly up-regulated in CR × CR larvae but not HR × HR larvae under heat that were also significantly up-regulated under control in HR × HR vs CR × CR larvae were identified as potentially front-loaded genes in HR × HR larvae. The same approach was repeated for determining back-loaded genes as well as front-loaded/back-loaded genes in CR × CR larvae.

To understand the functional implications of conserved gene modules identified in WGCNA, a categorical gene ontology (GO) enrichment analysis was performed using binary values (1 or 0) to indicate module membership in the WGCNA set followed by a Fisher’s exact test and false discovery rate correction. For heat-responsive DEGs by larval origin, signed log p-values using adaptive clustering of GO categories and a two-sided Mann-Whitney U-test was applied, followed by a false discovery rate correction. GO scripts can be found at https://github.com/z0on/GO_MWU. Heatmaps of hierarchically clustered GO terms were generated using the pheatmap package (Kolde, 2012) in R.

Discriminant analysis of principal components (DAPC) was performed to explore the relative changes in global expression between treatment in different populations from 2019 and 2021 using the R package adegenet (Jombart, 2008). Variance stabilized data (VSD) were used to create the model and the number of PCs was chosen to capture at least 80% of transcriptional variance. Distribution of samples grouped by origin and treatment was visualized in density plots.

## Results

### Temperatures and bleaching history at study sites

SST was always cooler at HR than CR (Fig. 1) during the data collection period, although bleaching occurred at both sites in 2014 and 2015 (Gintert et al., 2018; Manzello et al., 2019). Bleaching was observed at CR in 2018 and during gamete collection in 2019 at CR, but not at HR. CR gametes were collected from colonies that didn’t show visual bleaching. In 2021, bleaching was not observed at either site, in line with the cooler temperature patterns (Fig. 1). In total, from the start of the most recent mass global bleaching event in 2014 to the 2021 spawn, CR experienced four bleaching events, whereas HR only experienced two events. Notably, HR did not experience bleaching between 2015 and 2021.

### Larval survival under acute heat stress

To achieve a reasonable separation of survivorship among the different groups, the 36 °C acute temperature stress lasted for 96 hrs in 2019 and 141 hrs in 2021. The average survival rate at the end of each year’s experiment for the control vs. heat group was 77% vs. 47% and 86% vs. 33% respectively (Fig. 2). Exposure to 36 °C significantly increased larval mortality in both years, resulting in hazard ratios of 3.1 (*z* = 4.16, *p* < 0.001) and 28.7 (*z* = 4.11, *p* < 0.001) for heat-treated larvae from 2019 and 2021, respectively. Larval origin also had a significant effect on survival in both years. In 2019, the CR × CR cross experienced almost double the mortality risk (HR = 1.9, *z* = 2.18, *p* < 0.05) in comparison to the HR × HR cross, and in 2021 the CR × HR cross experienced more than 10 times the risk (HR = 11.8, *z* = 2.18, *p* < 0.001) of the HR × HR cross.

**Fig. 2.**
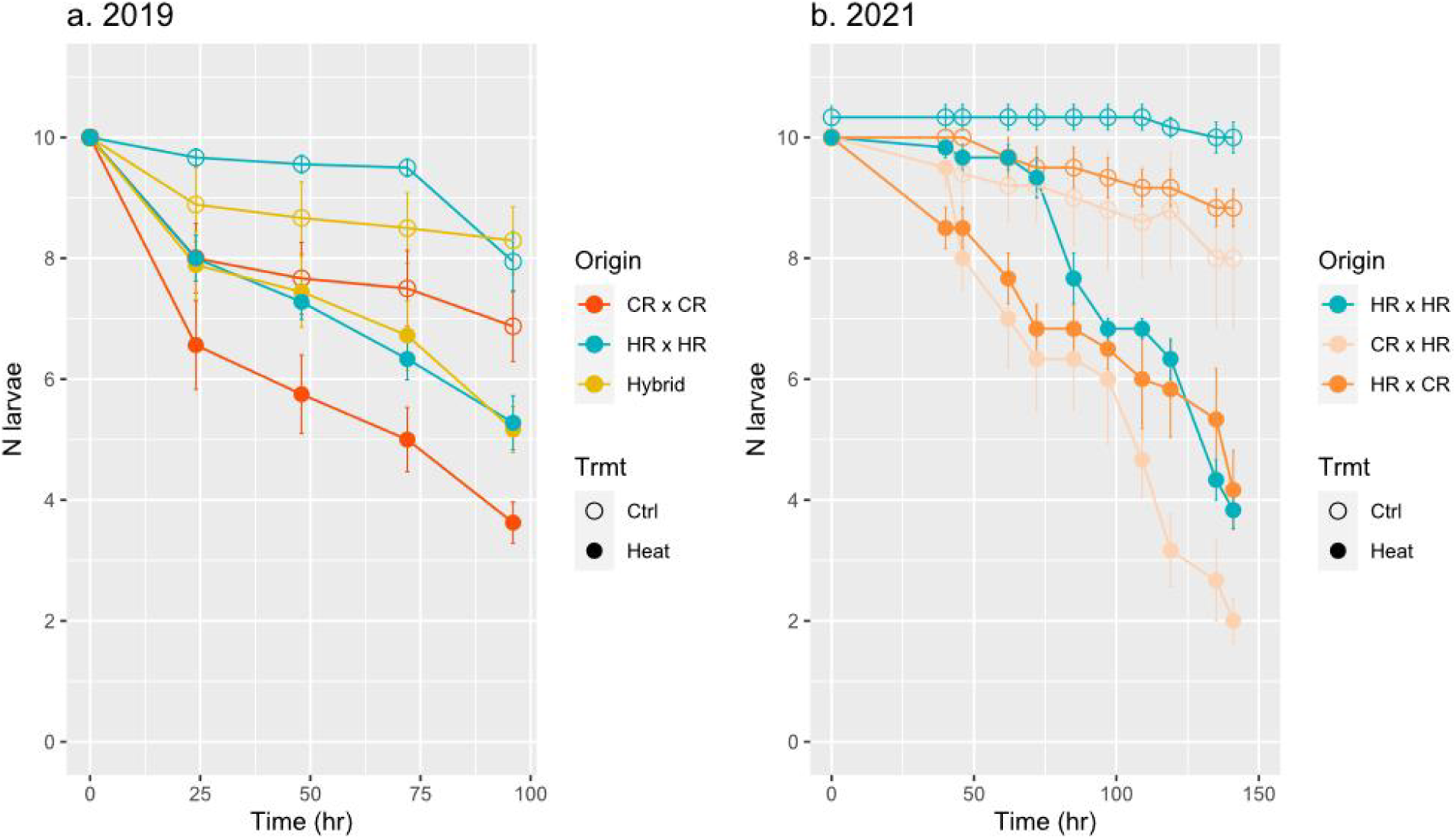
Number of surviving larvae (mean ± standard error of the mean [SEM]) across time in the acute temperature stress experiment for a) 2019 and b) 2021 cohorts. Survivorship was grouped by larval origin and treatment condition.

### Gamete (2019 and 2021) and larval ecophysiology (2021 only)

Total lipid content of gametes collected from CR and HR during 2019 spawning season did not differ between the two sites (Fig. 3b). However, HR gametes contained 2.2 times more wax esters (F = 6.27, df = 1, *p* < 0.05) and 1.5 times more phospholipids (F = 6.07, df = 1, *p* < 0.05) than CR gametes (Fig. 3a). No differences in triacylglycerol content by origin were apparent. Qualitatively, lipid content tended to be higher in the CR gametes in 2021 (Fig. 3b), but as only one colony spawned, formal significance was not evaluated. No differences were detected in total lipid content, lipid classes, or total soluble protein content of 2021 larvae between origin, treatment conditions or their interaction (Fig. S2, S3; Table S2). Protein and lipid data were unavailable for 2019 larvae exposed to the 32 °C moderate stress.

**Fig. 3.**
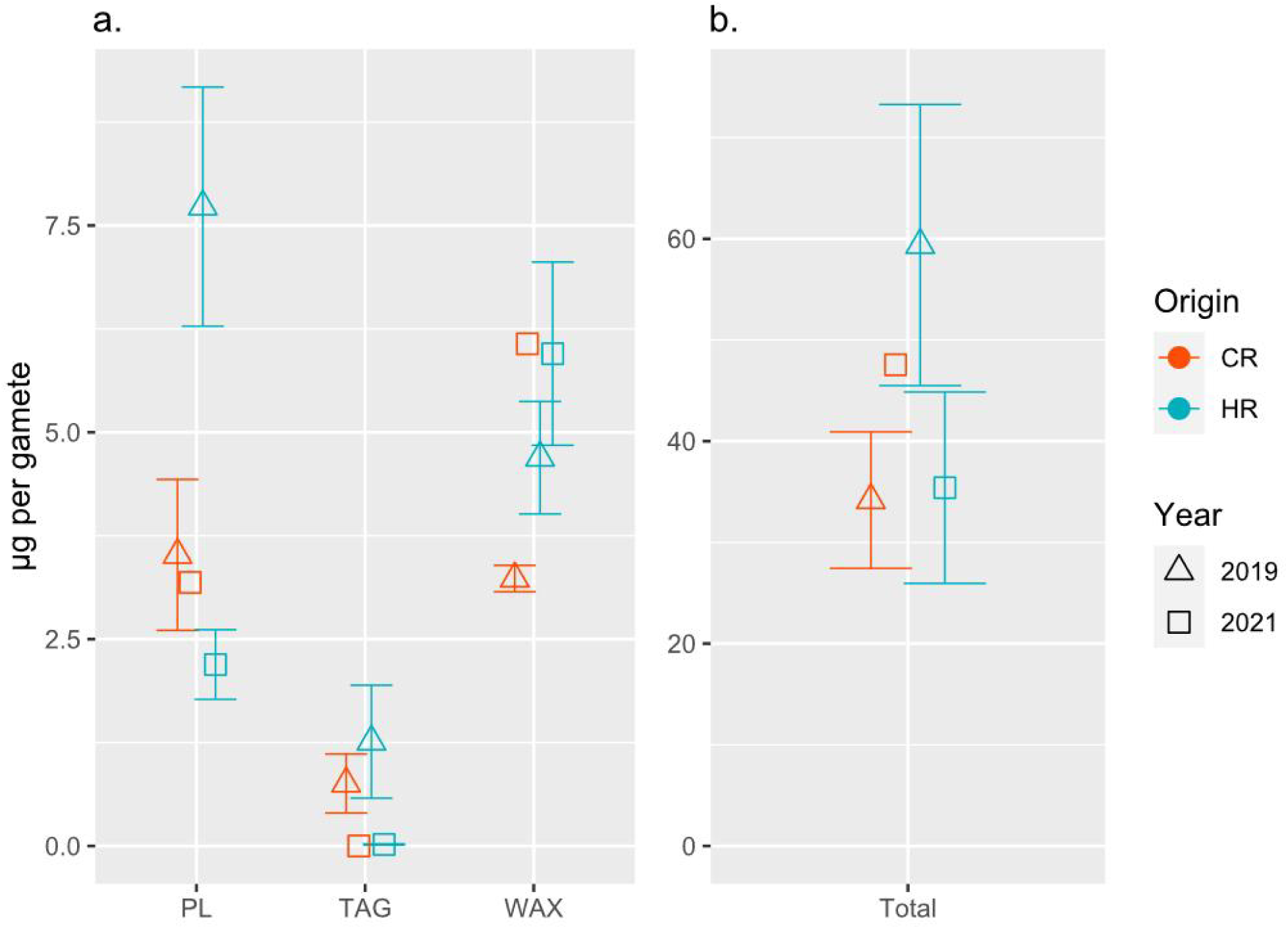
Concentration (mean ± SEM) of (a) different lipid classes (PL: phospholipid, TAG: triacylglycerol, WAX: wax ester) and (b) total lipids standardized by individual gamete bundle collected from Cheeca Rocks (n = 5) and Horseshoe Reef (n = 5) in 2019 and 2021. No replicates were available for CR gametes in 2021.

### Major drivers of transcriptional variation

Principal component analysis (PCA) on all high expression genes (count > 2 in 90% samples) within each dataset showed that larval origin was an important driver of transcriptional variance in addition to temperature treatment (Fig. 4). Samples clustered by larval origin along PC2 in the 2019 dataset, which accounted for 5.6% of the overall variance in expression (Fig. 4a). Putative hybrid samples generally clustered mid-way between HR × HR and CR × CR origin larvae. In the 2021 dataset, larval origin was the primary driver of expression variation, as cross types were clustered along the first PC which accounted for 10.4% of the overall variance (Fig. 4b). No significant differences in clustering were apparent between the reciprocal hybrids (CR × HR vs. HR × CR) but both were distinct from the HR × HR origin larvae. Clustering by temperature treatment was apparent along PC2 in the 2021 dataset, which explained 9.6% of the variance in overall expression (Fig. 4b), and along PC3 in the 2019 dataset, which explained 5.1% of transcriptional variance (Fig. S4).

**Fig. 4.**
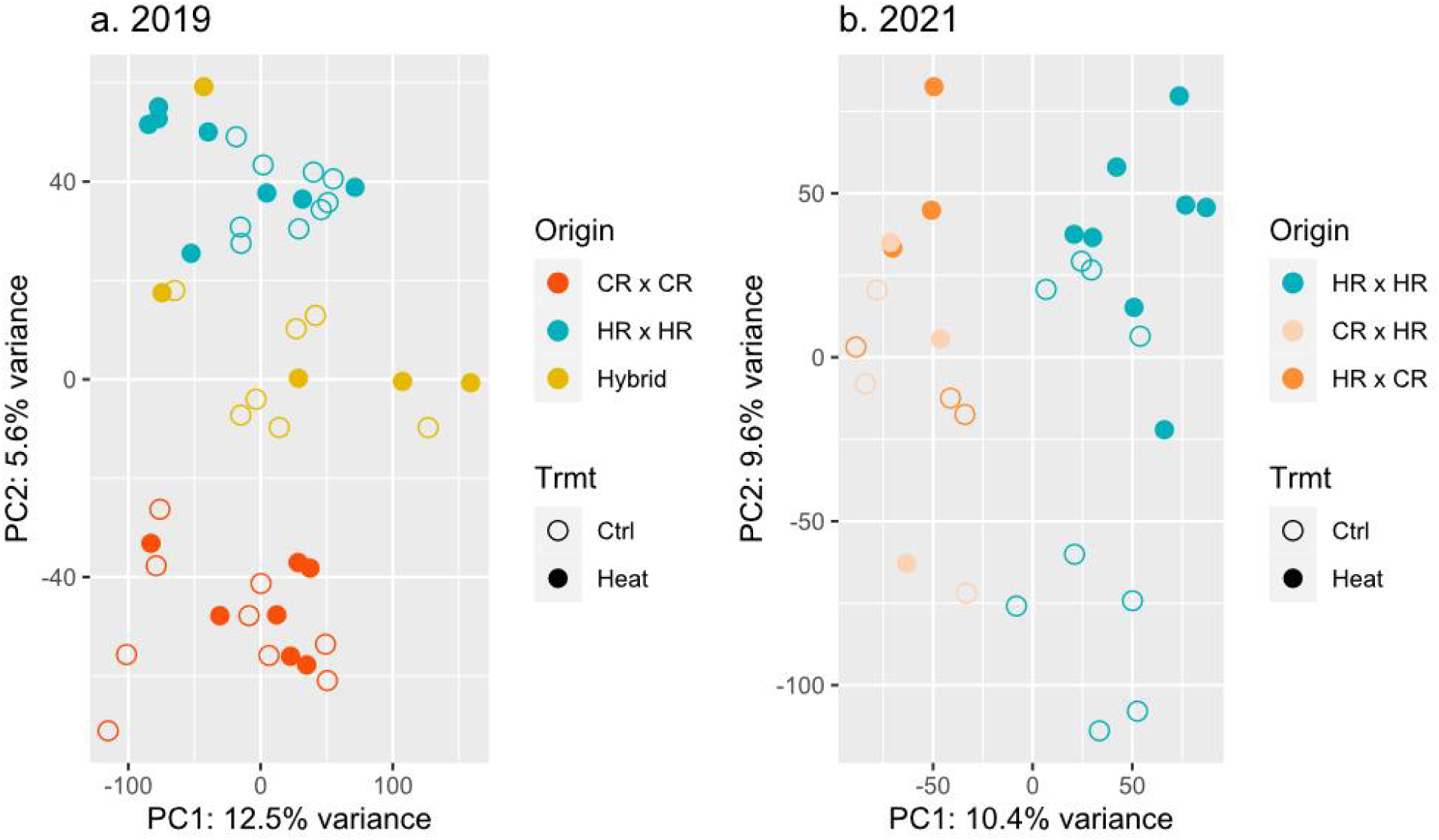
Principal component analysis (PCA) on *rlog*-transformed read counts in a) 2019 and b) 2021 larval datasets. Points are colored by origin and shaped by treatment. The percentage variance explained by each PC is reflected on the axis label.

### Conservation of expression networks and their functional significance

WGCNA was used to investigate whether and to what degree expression patterns were conserved between the two datasets and to further explore their relationships with experimental factors. Four modules, pink (n = 400 genes), purple (n = 268), magenta (n = 283), and red (n = 514), were highly correlated with origin in both years, although the directions of the correlations were not always conserved (Fig. 5). Specifically, genes within the pink and purple modules were strongly negatively correlated, or down-regulated, in 2019 CR × CR larvae (pink: Pearson’s r = - 0.87, *p_cor_* = 5e^-15^; purple: Pearson’s r = -0.74, *p_cor_* = 8e^-9^) and up-regulated in 2019 HR × HR larvae (pink: Pearson’s r = 0.81, *p_cor_*= 1e^-11^; purple: Pearson’s r = 0.69, *p_cor_* = 1e^-7^), while the opposite relationship was observed in the magenta (CR × CR: Pearson’s r = 0.71, *p_cor_* = 6e^-8^; HR × HR: Pearson’s r = -0.86, *p_cor_* = 7e^-14^) and red modules (CR × CR: Pearson’s r = 0.88, *p_cor_* = 2e^-15^; HR × HR: Pearson’s r = -0.76, *p_cor_* = 1e^-9^). In comparison, the magnitude and direction of module-trait correlations remained similar for the pink and red modules in 2021, with pink module genes again showing strong up-regulation in HR × HR larvae and red module genes showing strong down-regulation (pink: Pearson’s r = 0.98, *p_cor_* = 3e^-21^; red: Pearson’s r = -0.98, *p_cor_* = 3e^-21^). While there were still highly significant correlations observed for the purple and magenta modules with respect to origin, the direction of the association was completely reversed, with strong down-regulation of genes in the purple module (Pearson’s r = -0.98, *p_cor_* = 9e^-20^) and up-regulation of genes in the magenta module (Pearson’s r = 0.96, *p_cor_*= 5e^-17^). Modules significantly associated with treatment exhibited more moderate correlation coefficients (Pearson’s r range: ± 0.4 to ± 0.6), but their expression patterns were more strongly conserved across years. Genes in the black (n = 408 genes), greenyellow (n = 268), salmon (n = 160), and yellow (n = 574) modules were consistently up-regulated in heat-treated larvae whereas genes in the cyan (n = 152) module were consistently down-regulated in heat-treated larvae across years.

**Fig. 5.**
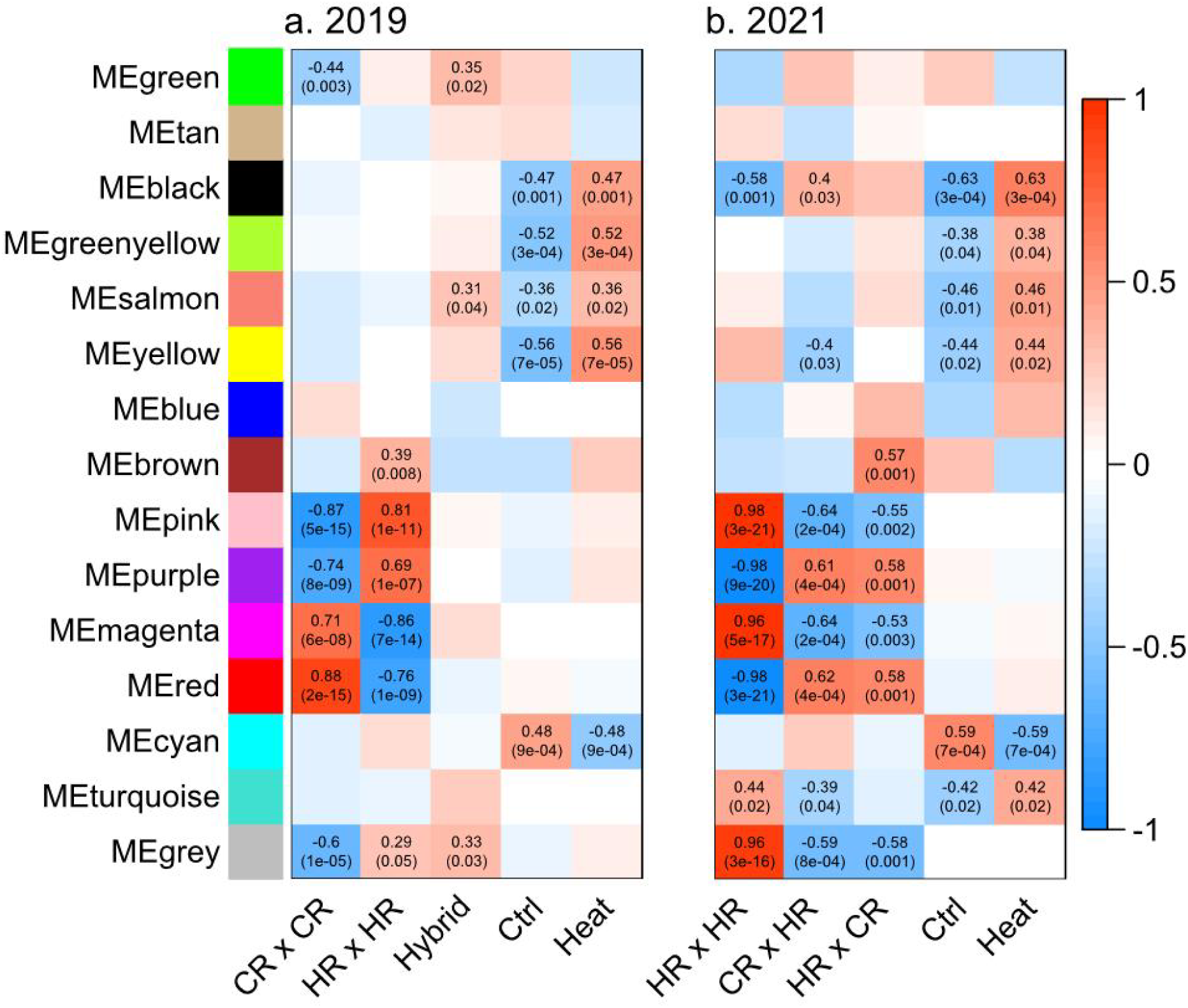
Weighted gene co-expression network analysis (WGCNA) module-trait relationships identified in a) 2019 and b) 2021 larval cohort. Correlation values range from 1 (red) to -1 (blue) and the associated *p* values were included in the parenthesis below for modules showing a significant trait association, with color of each block determined by strength and direction of the correlation between given module and trait.

Categorical functional enrichment analysis of genes assigned to the significant origin modules (pink, purple, red, magenta) revealed few significant GO terms, likely due to small module size. Two terms in the molecular function category (GO:0031683, G-protein beta/gamma-subunit complex binding; GO:0050780, dopamine receptor binding) and one term in the cellular components category (GO:0005834; GO:1905360, GTPase complex) were significantly enriched (*p* ≤ 0.05) among genes in the pink module (Table S3). GO analysis of treatment modules identified enrichment of genes associated with heat response pathways, such as immune response, defense response, and response to external stimulus in the black module which was up-regulated in response to heat treatment (*p* < 0.01, Fig. S5, Table S3). Whereas the cyan module, which was down-regulated in response to heat, showed enrichment of genes associated with amide and peptide metabolic and biosynthetic processes (*p* < 0.05, Fig. S6, Table S3). These processes were also enriched among genes in the salmon module, which was up-regulated under heat (Table S3). One term in the molecular function category (GO:0031210; GO:0050997, phosphatidylcholine binding) was significantly enriched (*p* = 0.05) among genes in the yellow module, and was also up-regulated under heat treatment (Table S3). No significant functional enrichments were detected for the greenyellow module.

### Origin-specific responses to thermal stress

DESeq2 analysis of the 2019 dataset showed that a total of 561 genes were up-regulated and 436 were down-regulated in heat-treated larvae relative to controls (Table S4). When further partitioned by origin, 133 heat-responsive genes were up-regulated and 144 were down-regulated in 2019 CR × CR larvae, whereas 376 were up-regulated and 158 were down-regulated in 2019 HR × HR larvae. The putative hybrid larvae up-regulated 12 genes and down-regulated 9 genes under heat and thus were excluded from the downstream functional enrichment analysis due to a limited number of differentially expressed genes (DEGs). Among these origin-specific heat-responsive genes, 28 (11 annotated) were front-loaded and 47 (31) were back-loaded in 2019 HR × HR larvae, while 82 (48) were front-loaded and 23 (14) were back-loaded in 2019 CR × CR larvae (Table S5). In the 2021 dataset, 706 heat-responsive genes were up-regulated and 953 were down-regulated in HR × HR larvae, 228 were up-regulated and 54 were down-regulated in CR × HR larvae, and 124 were up-regulated and 101 were down-regulated in HR × CR larvae.

Subsequent ontology analysis of these DEGs showed that biological processes including immune response, peptide hormone processing, defense response, and response to stimulus were enriched among heat-responsive up-regulated genes in 2019 CR × CR larvae (FDR < 0.01), while RNA metabolic process and macromolecule biosynthetic process were enriched among down-regulated genes (FDR < 0.01, Fig. 6a, Table S6). In the 2019 HR × HR larvae, phospholipid catabolic process was enriched among genes up-regulated in heat (FDR < 0.05), while microtubule-based movement/process and protein-DNA complex subunit organization were enriched among downregulated genes (FDR < 0.01, Fig. 6b, Table S7). Discriminant analysis of principal components (DAPC) for heat responsive genes revealed a greater transcriptional response in 2019 HR × HR larvae compared to the CR × CR larvae from the same year (Fig. 6c).

**Fig. 6.**
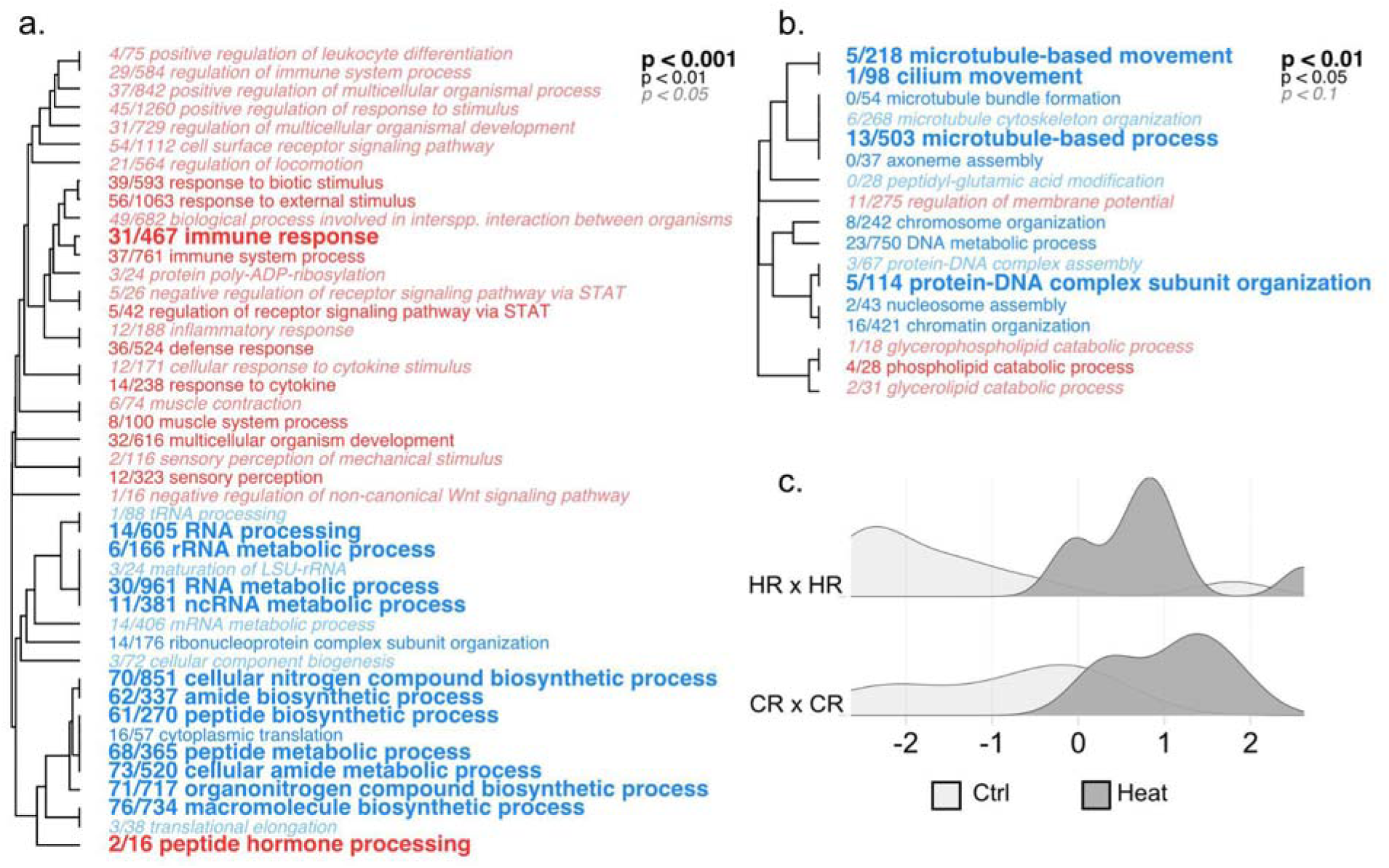
Hierarchical clustering of ontology terms enriched by genes up-regulated (red) or down-regulated (blue) in 2019 heat-treated (a) CR × CR larvae and (b) HR × HR larvae compared to their respective untreated control, summarized by biological process (BP). Font size indicates level of statistical significance (FDR-corrected). Term names are preceded by fractions indicating the number of individual genes within each term differentially regulated with respect to treatment (unadjusted *p* < 0.05). (c) Density plots showing distribution of global expression across samples from the two origins along the temperature responsive axis (linear discriminant 2 [LD2], Fig. S7) based on discriminant analysis of principal components (DAPC) performed on variance stabilized data (VSD) grouped by treatment and origin.

In the 2021 cohort, genes up-regulated in response to heat in HR × HR larvae were enriched for biological processes including DNA metabolic process, organelle localization, and cellular response to DNA damage stimulus (FDR < 0.01, Fig. S8, Table S8). Processes enriched among down-regulated genes included a suite of metabolic processes of small and large molecules, such as organic acids, fatty acids, and lipids (FDR < 0.01). For the two hybrid crosses, upregulated genes were enriched for NF-κB signaling regulation, immune and defense response, as well as regulation of cell death (FDR < 0.05, Fig. S9, S10, Table S9, S10). Downregulated genes were enriched for lipids and protein metabolic/catabolic processes (FDR < 0.05). DAPC for heat responsive genes identified a greater response in HR × CR larvae compared to the HR × HR and CR × HR larvae from the same year (Fig. S11).

## Discussion

Adult *O. faveolata* from Cheeca Rocks exhibit elevated thermal tolerance in response to natural bleaching events (Gintert et al., 2018; Manzello et al., 2019) suggesting that they have acclimatized or adapted to naturally higher and more variable temperatures characteristic of inshore reef sites in the Florida Keys (Kenkel et al., 2015; Kenkel & Matz, 2016). Contrary to the current paradigm of inherited and/or enhanced thermal tolerance in adults experiencing more extreme thermal regimes (Dixon et al., 2015; Putnam & Gates, 2015; Strader & Quigley, 2022), we found that offspring of these more tolerant inshore colonies were more susceptible to thermal stress, exhibiting reduced survival and stronger expression signatures of a stress response in comparison to larvae from offshore colonies (Fig. 2, 6). The observed total lipid data suggest robust bleaching resistance in adults may come at the cost of reproductive investment (Fig. 3), although patterns are inconsistent across years. These findings represent an important counter-example to the rationale underpinning selective breeding approaches (Drury et al., 2022): that tolerant parents can be counted on to produce tolerant offspring.

### Impaired larval performance may result from reduced reproductive investment

Reef origin (or cross type) and temperature treatment played important roles in driving physiological traits in *O. faveolata* larvae. As expected, larvae were more likely to die under heat treatment than in the ambient control (Fig. 2), but the origin response was unexpected. Horseshoe Reef (HR) purebred larvae appeared to be the best performers in both years, while Cheeca Rocks (CR) purebred larvae in 2019 and 2021 CR × HR hybrids experienced significantly higher mortality in comparison (Fig. 2). Previous studies conducted on *Acropora millepora* and *Acropora tenuis* from the Great Barrier Reef showed that larval survival under a similar level of acute thermal stress (35.5 °C) was enhanced in offspring of parents from warmer source reefs (Dixon et al., 2015; Strader & Quigley, 2022). Yet, in our study system, the HR × HR cross, produced by the less tolerant parents sourced from a cooler reef environment, repeatedly outperformed the CR cross produced by more tolerant parents from a warmer reef environment (Gintert et al., 2018; Manzello et al., 2019). This suggests that the host genetic contribution to thermal tolerance may be minimal or overpowered by other factors such as recent and concurrent heat stress, or maternal provisioning.

Prior thermal stress and bleaching has been linked to negative reproductive outcomes, including both fecundity and gamete quality, in multiple coral species (Jones & Berkelmans, 2011; Szmant & Gassman, 1990; Ward et al., 2002). Although the severity of reproductive impacts is thought to be related to the severity of bleaching and rate of recovery, resistant/resilient colonies do not necessarily exhibit latent effects (Leinbach et al., 2021; Szmant, 1991). Notably, the HR source population suffered from both a reduced number of spawning colonies and total gametes released as a result of the back-to-back bleaching events in 2014 and 2015 (Fisch et al., 2019). Previous research consistently demonstrates that adult colonies from nearshore reef environments display higher resistance and resilience to thermal stress compared to those from offshore reefs (Gintert et al., 2018; Manzello et al., 2019). Based on these findings, we anticipated that the offspring of these resilient adults would also exhibit enhanced thermal tolerance, in line with previous studies. At the time of spawning collections in 2019, temperature at CR had surpassed the local bleaching threshold and a majority of the colonies showed some degree of paling (i.e., onset of bleaching), while HR experienced cooler temperatures and all the colonies appeared fully pigmented (Fig. 1). Recently, (Leinbach et al., 2021) showed that *Acropora hyacinthus* which resisted bleaching maintained higher reproductive capacity than recovered coral. Although we only collected gametes from CR colonies without apparent signs of physiological stress in 2019 (i.e. thermally resistant adults), it is likely that the energetic state of individual colonies was already compromised due to the heat stress, which may have impacted maternal provisioning.

Scleractinian coral gametes are largely composed of lipids with wax esters, phospholipids and triacylglycerols being the most abundant classes (Figueiredo et al., 2012). Mean total lipids were higher for 2019 HR gametes compared to CR gametes (Fig. 3b). Lipid class analyses, of the 2019 gametes, showed CR gametes contained less wax esters and phospholipids than HR gametes (Fig. 3a). Wax esters take longer to metabolize, supporting their role in long-term energy storage, and are important for larval development (Lee et al., 2006; Richmond, 1987; Rivest et al., 2017), indicating 2019 HR gametes had greater energy reserves available for dispersal and settlement. Additionally, greater wax ester stores lower gamete/larval density, possibly contributing to extended dispersal (Richmond, 1987). Phospholipids were significantly different across sites in 2019, but this lipid class encompasses many structural lipid compounds, thus muddling the implications. While triacylglycerols can be rapidly hydrolyzed and likely support immediate energetic needs (Figueiredo et al., 2012; Sewell, 2005), no significant differences were detected across sites in either 2019 or 2021. Similar lipid class patterns were not observed in the 2021 cohort, possibly due to lack of replication in CR gametes (Fig. 3a). The general deficiency of 2019 CR gametes in all these classes suggests that bleaching tolerance and resilience of CR adults may come at the cost of reproductive investment, potentially contributing to reduced performance of CR larvae.

Further supporting a compromised reproductive output of nearshore coral populations was the absence of mass spawning on CR in the summer of 2021, despite no observed bleaching (Fig. 1), which limited our ability to re-assess performance of CR × CR larvae and additional hybrid crosses. This implies that reproductive capacity of CR adults (likely among other marginal nearshore populations) could be jeopardized by the persistent latent effects of accumulated stress. As thermal anomalies increase in magnitude and frequency in the Florida Reef Tract (Manzello, 2015), it is important to consider any latent effects beyond visible stress responses that could affect the next generation and ultimately the persistence of reef communities.

### Stress tolerance rather than a front-loaded stress response is associated with enhanced larval survival

In addition to origin-specific physiological differences, larvae originating from different reef sites mounted different transcriptional responses after four days of 32 °C heat challenge. In the 2019 CR purebred larvae, we observed functional enrichment in a wide range of stress responses (e.g., immunity, defense, inflammation) among the significantly upregulated genes, whereas metabolic and biosynthetic processes (e.g., various RNA molecules, peptide, cellular nitrogen compounds) were enriched among downregulated genes (Fig. 6a). Similarly, underperforming CR × HR larvae in 2021 showed pronounced upregulation of defense and immune response pathways and downregulation of metabolic processes (Fig. S9). Upregulation of stress response genes and concomitant downregulation of growth related processes, such as rRNA metabolism, is a hallmark of the environmental stress response (López-Maury et al., 2008). In contrast, fewer enriched GO categories were identified among the heat-responsive DEGs in the 2019 and 2021 HR purebred crosses (Fig. 6b, S7), and more interestingly, the enrichments did not highlight stress response pathways like their CR × CR (2019) or hybrid (2021) counterparts, but cellular homeostatic processes instead (Fig. 6b).

The lack of an apparent stress response in HR purebreds does not appear to be due to an inability to detect differential expression as a result of transcriptional front-loading, or higher baselines expression of stress response genes (Barshis et al., 2013) We tested for front-loading and found that comparatively fewer genes were front-loaded in HR × HR larvae and none of those were annotated as stress response genes (Table S5). Among the annotated genes that were front-loaded in CR × CR larvae, two were related to protein ubiquitination (Ube2g1 and ZNF598, Table S4), which may indicate constitutive expression of stress response pathways. Additionally, HR purebreds had a more robust expression response to thermal stress, exhibiting lower baseline expression, but achieving the same magnitude of overall transcriptional plasticity in response to thermal stress as 2019 CR × CR larvae (Fig. 6c). In 2021 HR × HR and CR × HR larvae showed similarly elevated levels of baseline expression in comparison to HR × CR hybrid (Fig. S11). Taken together, this suggests the 2019 HR × HR larvae may be more resistant to thermal stress not because they were pre-conditioned for stressful conditions, but because they were able to strongly and rapidly acclimate their physiology, possibly as a result of having more energy reserves to devote towards a stress response (Fig. 3). Such a robust response may also be followed by a rapid return to baseline expression, or transcriptomic resilience (Rivera et al., 2021), when stress abates, although additional time-course data are needed to test this hypothesis.

### Conserved transcriptomic signatures of population origin and response to treatment

In addition to survival and gamete lipid content, global gene expression profiles of aposymbiotic *O. faveolata* larvae also revealed a strong signature of reef origin. The 2019 samples were organized into 3 distinct clusters based on origin along PC2, while origin was the predominant driver of clustering in the 2021 samples (Fig. 4). Treatment appeared to be a weaker driver in both datasets, clustering samples along PC3 in 2019 and PC2 in 2021 (Fig. S4, 4b). Despite a difference in developmental age of the two larval cohorts at the time of sampling (7 vs. 9 days post fertilization for 2019 vs. 2021), a WGCNA meta-analysis identified highly conserved gene modules significantly correlated with larval origin although the magnitude and direction of expression patterns was not always conserved (Fig. 5). Corals exhibit waves of transcription during early development consistent with zygotic genome activation and degradation of maternal transcripts (Chille et al., 2021; Hayward et al., 2011). Differences in the magnitude and direction of select modules between the 2019 and 2021 datasets (purple and magenta, Fig. 5) may reflect these temporal transcriptional waves, although a more thorough time course is needed to test this hypothesis. Nevertheless, evidence of such strong module preservation implies the existence of a core group of origin-specific genes that have a lasting effect throughout the organisms’ development and these modules may be linked to baseline differences in larval physiology. However, little information on the functional implications of these gene sets was retrieved from the GO enrichment analysis, which may be attributable to small module size and/or an insufficient number of annotated genes. This could be a worthwhile investigation for future studies as annotations improve and better enrichment methods become available.

WGCNA meta-analysis also identified highly conserved gene modules that were significantly associated with temperature treatment. Similar to the origin-specific modules, the majority of these treatment modules lacked significant functional enrichment. For the two modules that did have sufficient enrichment, biological processes including immune response, defense response, and response to external stimulus were enriched in genes associated with the black module, which were up-regulated in heat (Fig. S5), while amide and peptide metabolic and biosynthetic processes were enriched in genes associated with the cyan module that were downregulated in heat (Fig. S6). Therefore, upregulation of genes involved in stress response pathways and downregulation of metabolic processes related to growth and development in heat-treated larvae aligns with the cellular stress and cellular homeostasis response profiles identified in corals (and other marine and terrestrial organisms) during short- to medium-term physiological stress (Kenkel et al., 2014; Kültz, 2005).

### Implications for adaptive management

The documented resilience of adult corals to recurrent heat stress and bleaching in environments with high and variable temperatures may come with highly consequential trade-offs. In this study, we show that larvae from a site that routinely experiences and recovers from heat-induced bleaching, significantly underperformed relative to larvae from a cooler, less variable reef site. The unexpected outcome that more thermally tolerant *O. faveolata* produced poorer quality offspring challenges the prevailing paradigm that breeding vulnerable populations with thermally tolerant individuals can contribute to genetic rescue (Bay et al., 2017), which also serves as the theoretical justification for selective breeding approaches (Drury et al., 2022). Moreover, natural adaptive capacity is likely already impaired as CR × CR larvae and CR × HR hybrids exhibited greater mortality risk even under ambient conditions (Fig. 2). These findings also align with the long-term pattern of recruitment failure in Caribbean coral (Hughes & Tanner, 2000; Williams et al., 2008)

## Data availability

Raw sequence data can be found under NCBI BioProject PRJNA981197. Scripts and input files are available at DOI: 10.5281/zenodo.8025715.

## Acknowledgements

This project was supported by the NOAA OAR ‘Omics Program through an award to D. Manzello/I. Enochs and start-up funds from the University of Southern California to C. Kenkel. We thank A. Mayfield for his field support with the 2019 experiments. We thank the Baker Lab and the Traylor-Knowles Lab at the University of Miami RSMAS for their technical support with the 2021 experiments.

