## Supplemental materials for "Performance of *Orbicella faveolata* larval cohorts does not align with previously observed thermal tolerance of adult source populations"

**Supplementary Materials**

*Orbicella faveolata* reference transcriptome assembly and annotation

Replicate fragments from seven coral colonies representing three inshore and four offshore host genotypes (Manzello et al. 2019) were collected in July 2017 and subjected to a short and moderate-duration thermal stress experiment (Table S1). Briefly, samples were acclimated for seven days to control conditions (30°C) in Experimental Reef Lab (ERL) aquaria followed by a 7-day temperature ramp for fragments assigned to thermal stress treatments to reach the moderate (32°C) and short (33°C) targets. Fragments in the short-duration treatment were sacrificed by snap-freezing in liquid nitrogen after 5 days at the target temperature and after 31 days in the moderate-duration treatment. Total RNA was extracted using the RNAqueous kit (AM1912, Life Technologies) and sent on dry ice to the Duke Center for Genomic and Computational Biology for library preparation and sequencing. Libraries were sequenced in a 75 bp PE run on the NextSeq500 High Output flow cell returning a total of 574 M raw reads. Adapters and low-quality reads were removed in Trimmomatic v0.36 (phred33, quality score > 20, 4 bp sliding window; Bolger, Lohse, and Usadel 2014) yielding a total of 498 M high quality reads used for a *de novo* assembly in Trinity v2.5.1 (Grabherr et al. 2011), yielding a total of 491,454 contigs (N50 = 1,192) in the metatranscriptome. The metatranscriptome was filtered using BLASTx (e-value < 1e^−5^) searches against two coral host proteomes, and BLASTn (e-value < 1e^−5^) searches against four Symbiodiniaceae genomes and transcriptomes (corals: *Acropora digitifera,* (Shinzato et al. 2011); *Orbicella faveolata*, (Prada et al. 2016); Symbiodiniaceae: *Symbiodinium microadriaticum,* (Aranda et al. 2016; Bayer et al. 2012), *Breviolum minutum* (Shoguchi et al. 2013; Bayer et al. 2012), *Cladocopium goreaui*, (Liu et al. 2018), *Durusdinium trenchii*, (Dougan et al. 2022)). Transcripts were sorted based on their lowest e-value and highest bit score, retaining only transcripts with best hits to host references. The results from a GenBank nr (non-redundant) database (Sayers et al. 2019) search (e-value < 1e^−4^) were used to identify contigs matching metazoan proteomes to corroborate candidate host transcripts. Transcripts with no hits to the nr database were also retained if they matched the coral databases. A total of 87,440 contigs (N50 = 2,228, mean GC content: 42%) were assigned to Orbicella faveolata. Host transcripts were annotated using BLASTx and BLASTp searches against the uniprot database (e-value cut-off = 1e^−4^, Bateman et al. 2017) using the Trinotate v3.1.1 pipeline (Bryant et al. 2017). Comparison against the Benchmarking Universal Single-Copy Ortholog set (BUSCO) revealed that 91% of the core metazoan orthologs were represented (Simão et al. 2015; Nishimura, Hara, and Kuraku 2017).

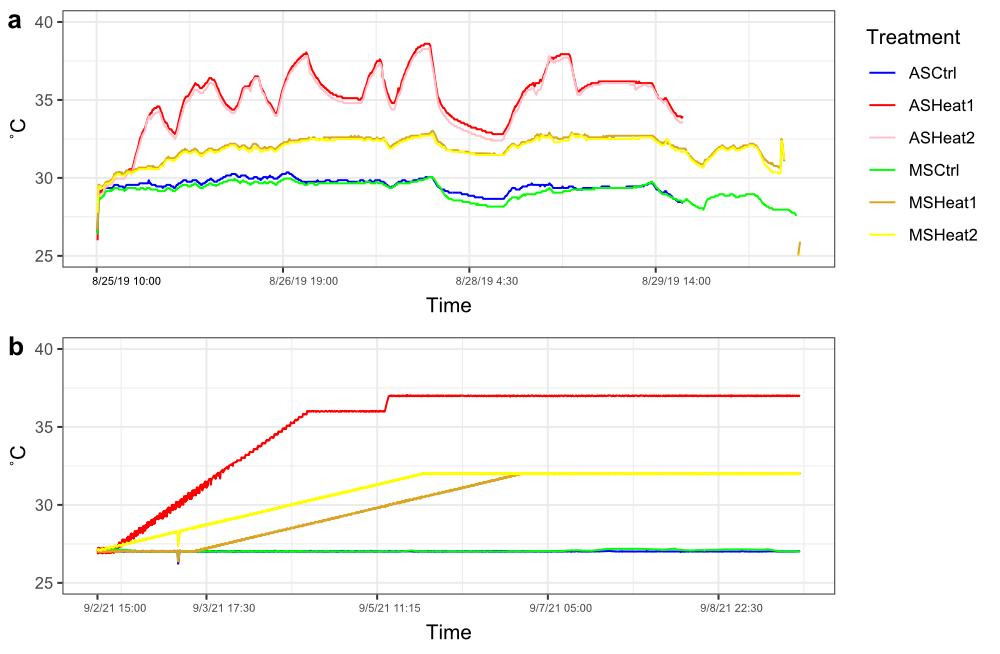

Fig. S1 Thermal profile of the 36 °C acute stress (AS) and 32 °C moderate stress (MS) experiments conducted in a) 2019 and b) 2021. Note that heat ramp for 2021 CR x HR and HR x CR larvae (MSHeat 1) started on 9/3/2021 15:00 due to the 24-hr developmental delay.

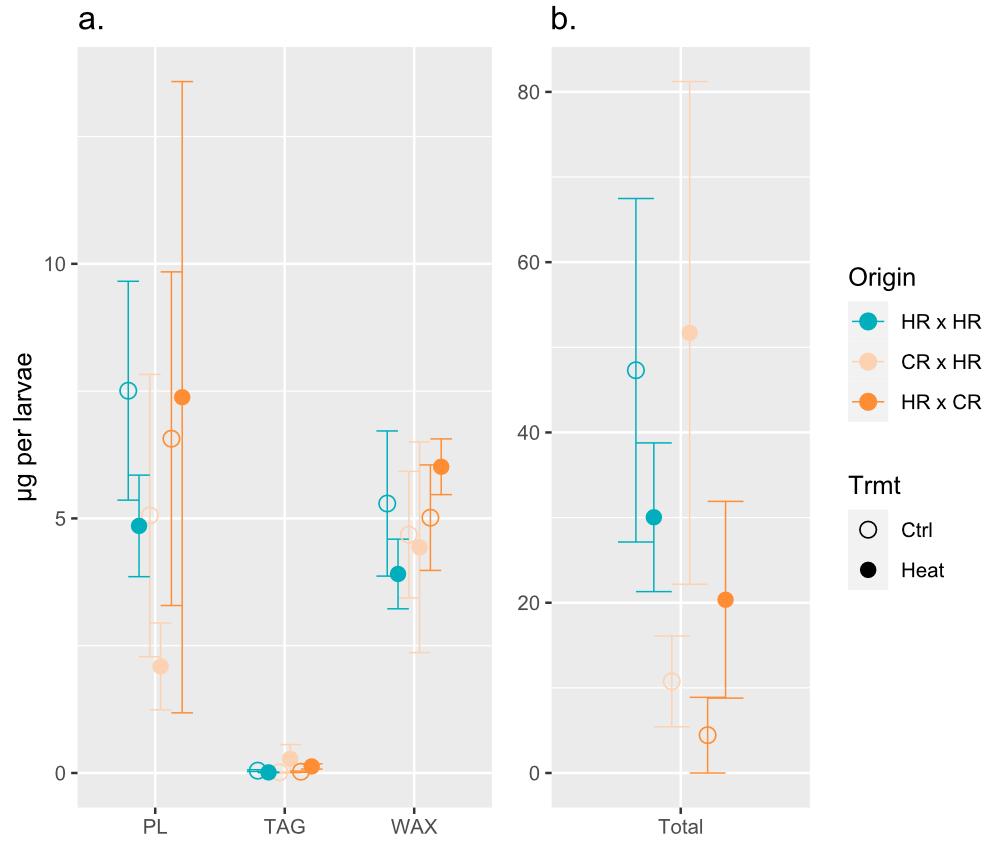

Fig. S2 Concentration (mean ± standard error of the mean [SEM]) of (a) different lipid classes (PL: phospholipid, TAG: triacylglycerol, WAX: wax ester) and (b) total lipids standardized by individual larvae in 2021. Points are colored by origin and shaped by treatment.

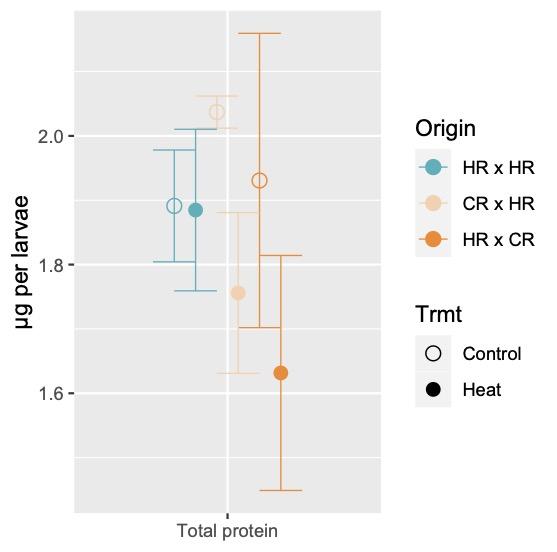

Fig. S3 Concentration (mean ± SEM) of total host soluble protein standardized by individual larvae in 2021. Points are colored by origin and shaped by treatment.

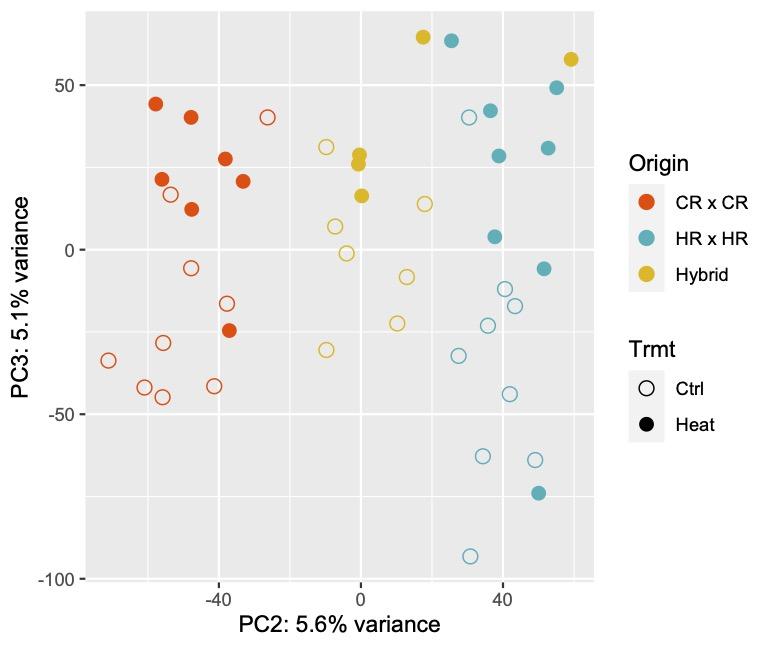

Fig. S4 Principal component analysis (PCA) on *rlog*-transformed read counts in 2019 larval dataset displaying PC2 and PC3. Points are colored by origin and shaped by treatment. The percentage variance explained by each PC is reflected on the axis label.

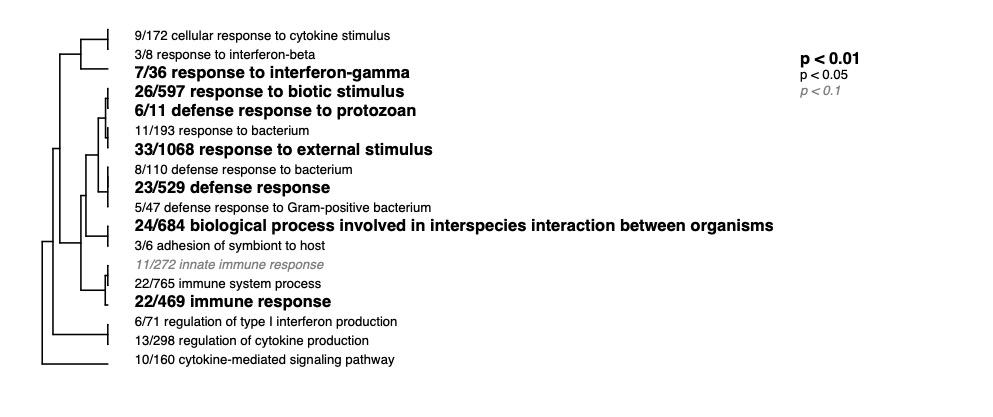

Fig. S5 Hierarchical clustering of ontology terms enriched by genes included in the back module determined by WGCNA, summarized by biological process (BP).

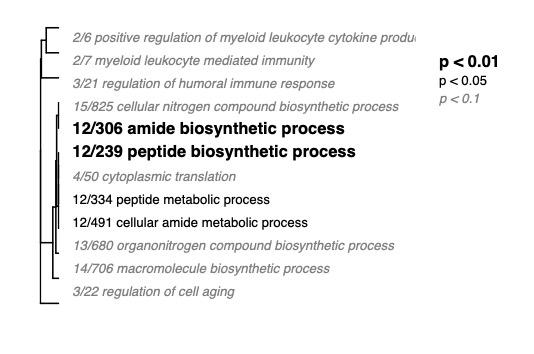

Fig. S6 Hierarchical clustering of ontology terms enriched by genes included in the cyan module determined by WGCNA, summarized by biological process (BP).

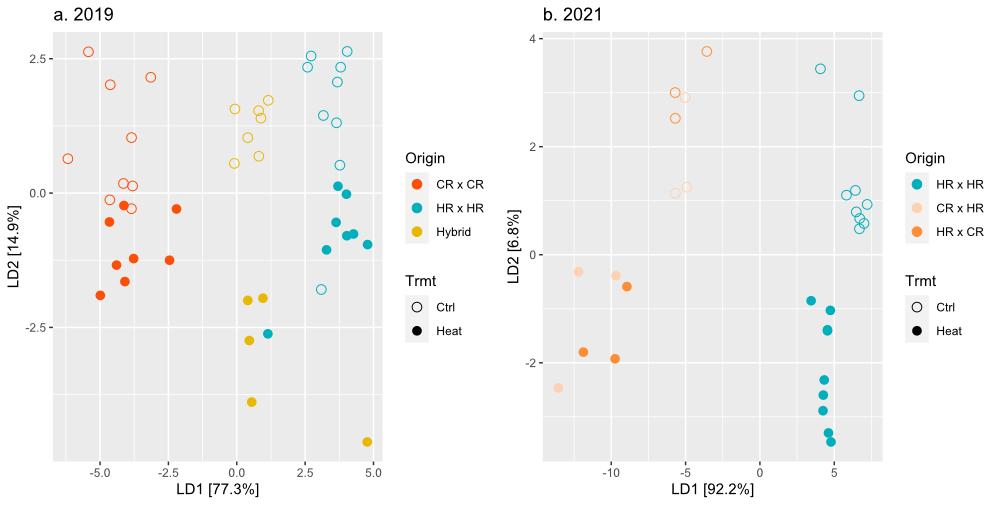

Fig. S7 Discriminant analysis of principal components (DAPC) on variance stabilized data (VSD) in a) 2019 and b) 2021 larval datasets. Points are colored by origin and shaped by treatment. The percentage variance explained by each linear discriminant (LD) is reflected on the axis label.

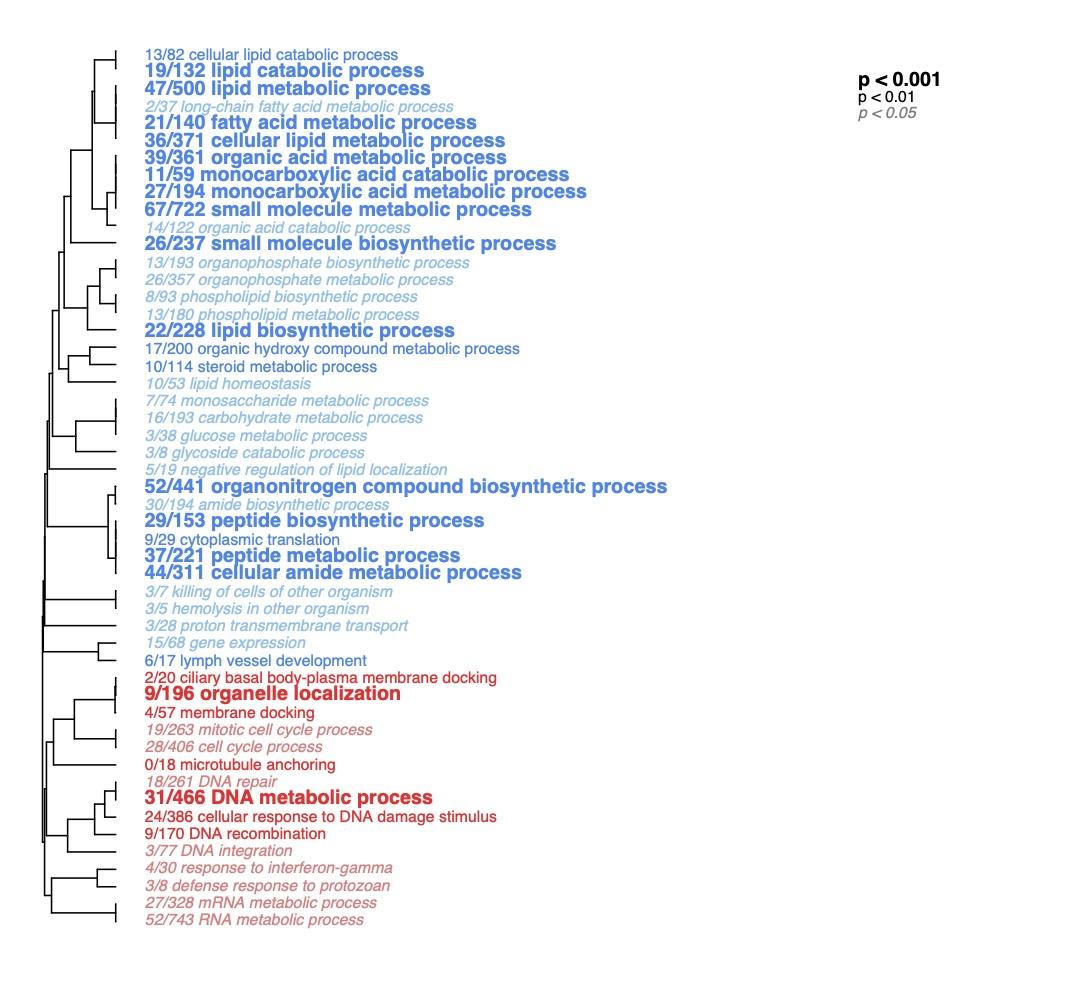

Fig. S8 Hierarchical clustering of ontology terms enriched by genes up-regulated (red) or down-regulated (blue) in heat-treated 2021 HR x HR larvae compared to their untreated control, summarized by biological process (BP). Font size indicates level of statistical significance (false discovery rate-corrected [FDR]). Term names are preceded by fractions indicating the number of individual genes within each term differentially regulated with respect to treatment (unadjusted *p* < 0.05).

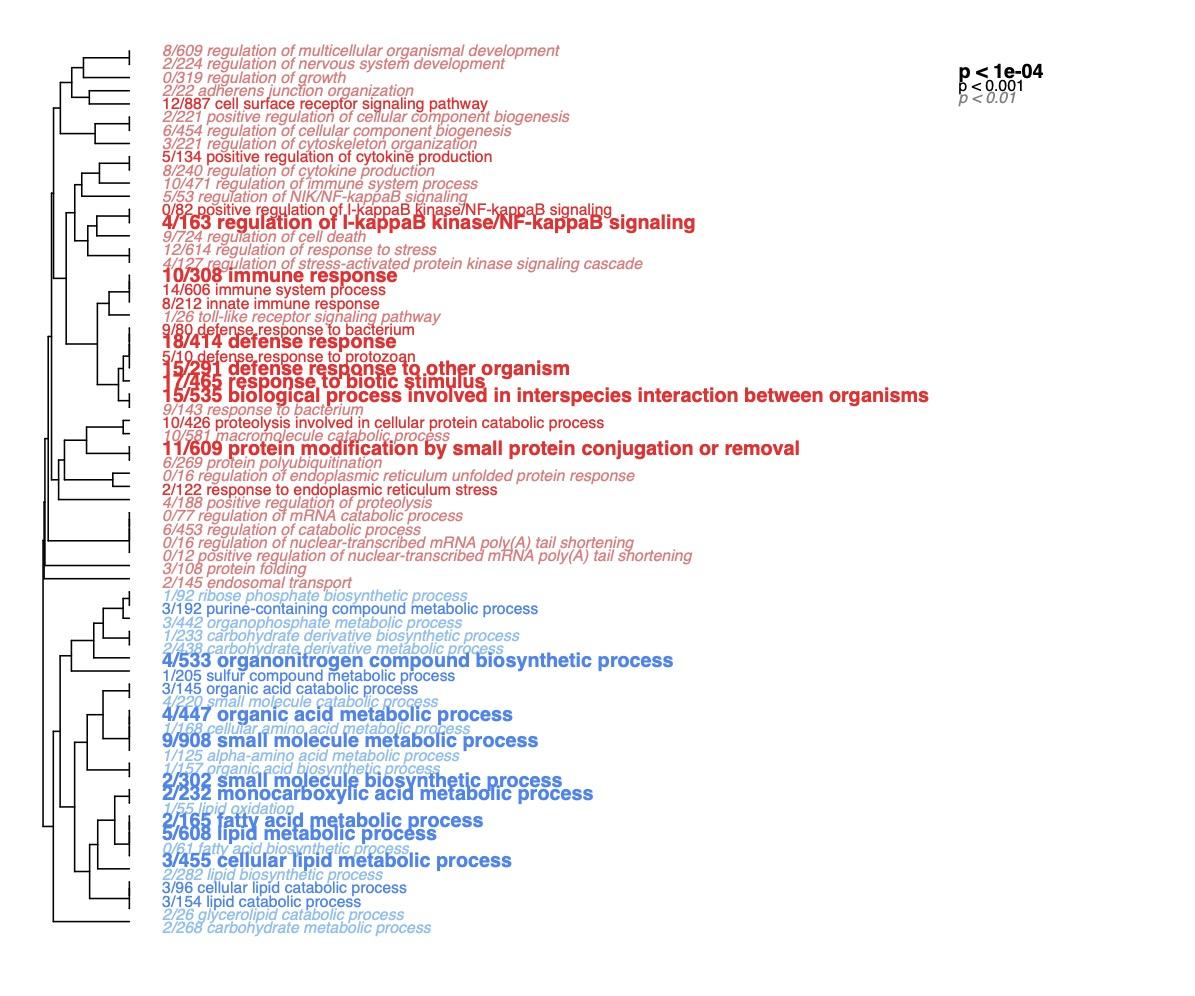

Fig. S9 Hierarchical clustering of ontology terms enriched by genes up-regulated (red) or down-regulated (blue) in heat-treated 2021 CR x HR larvae compared to their untreated control, summarized by biological process (BP). Font size indicates level of statistical significance (FDR-corrected). Term names are preceded by fractions indicating the number of individual genes within each term differentially regulated with respect to treatment (unadjusted *p* < 0.05).

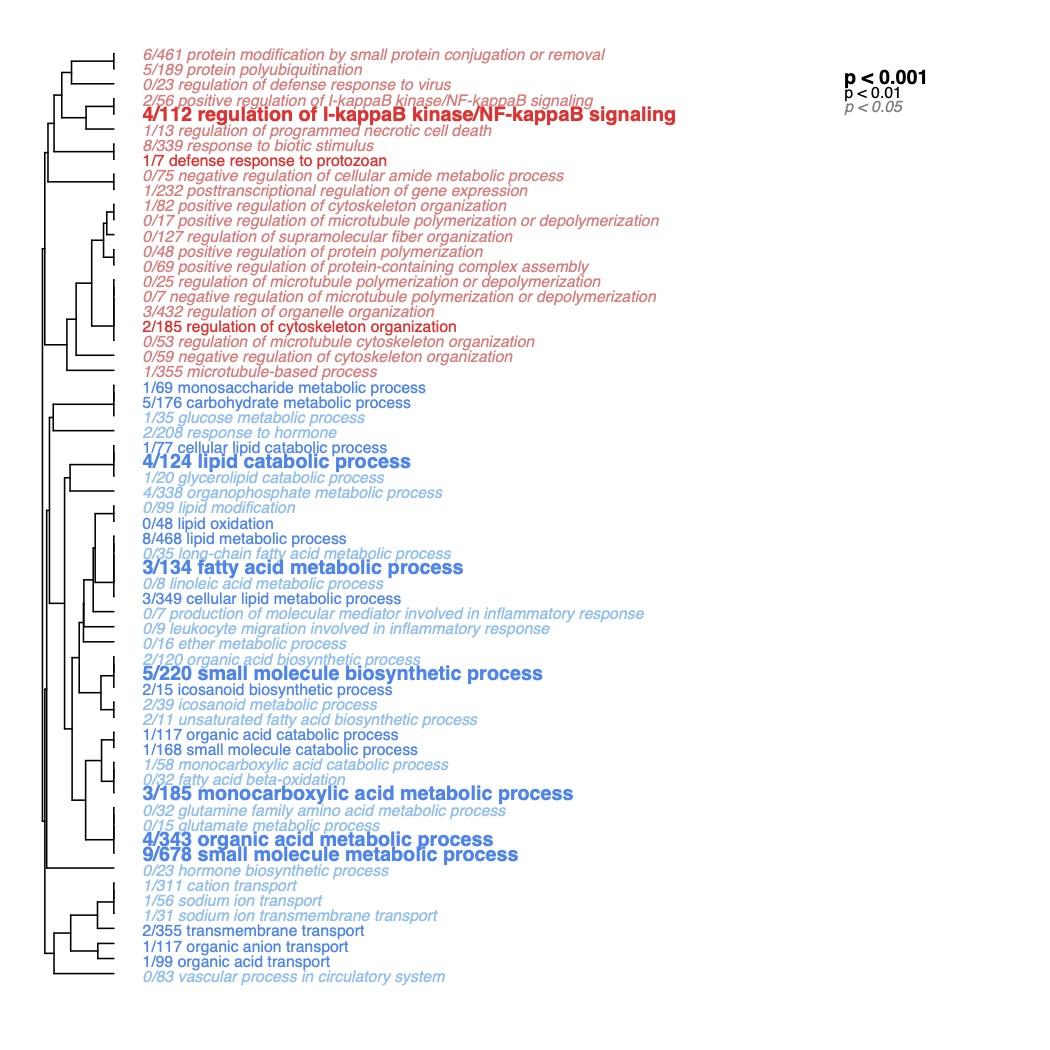

Fig. S10 Hierarchical clustering of ontology terms enriched by genes up-regulated (red) or down-regulated (blue) in heat-treated 2021 HR x CR larvae compared to their untreated control, summarized by biological process (BP). Font size indicates level of statistical significance (FDR-corrected). Term names are preceded by fractions indicating the number of individual genes within each term differentially regulated with respect to treatment (unadjusted *p* < 0.05).

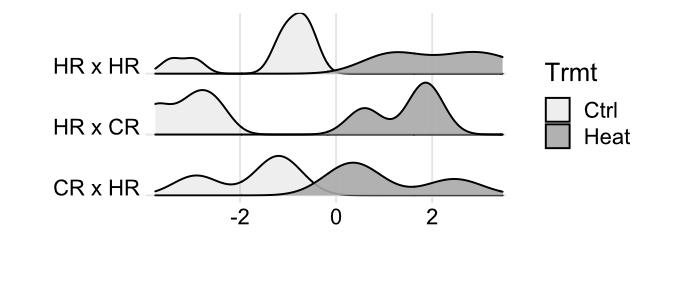

Fig. S11 Density plots showing distribution of global expression across samples from the three origins along the temperature responsive axis (LD2, Fig. S7) based on discriminant analysis of principal components (DAPC) performed on variance stabilized data (VSD) grouped by treatment and origin.
