## Supplemental Table 1 for "Performance of *Orbicella faveolata* larval cohorts does not align with previously observed thermal tolerance of adult source populations"

**Table S1 (adapted from Aguilar et al., *in prep*).** Experiment samples distribution. GPS coordinates for the collection sites are as follows: UKI1(24.89742°N, 80.61573°W), UKI2 (24.95375°N, 80.54806°W), UKO2 (24.9465°N, 80.50205°W).

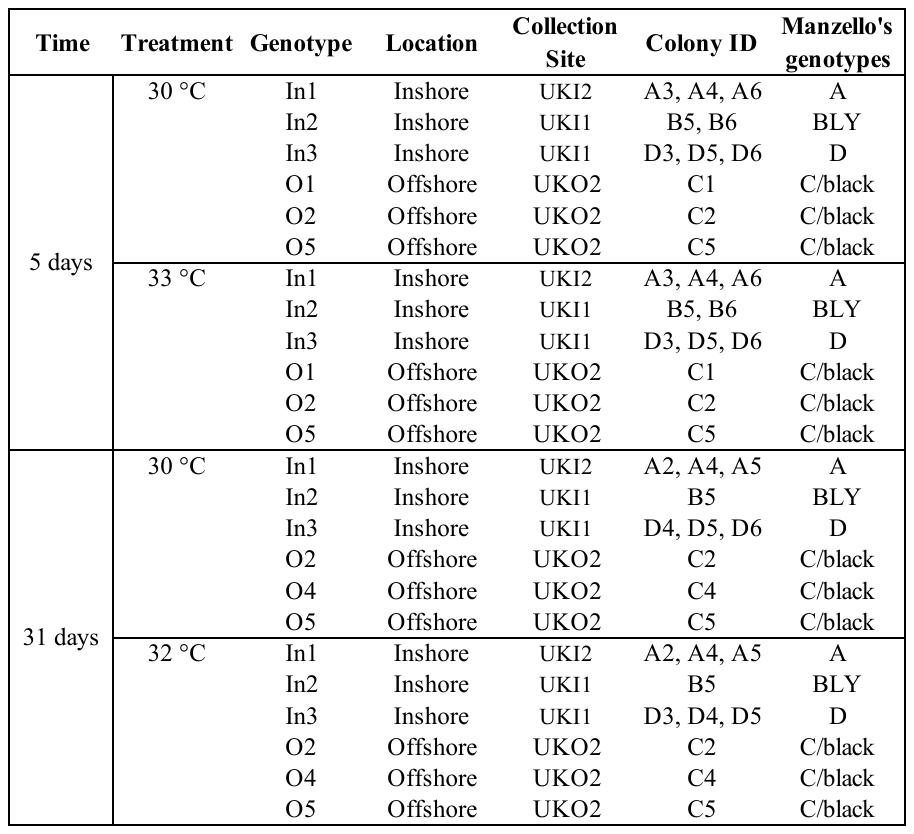
